# Cellpose3: one-click image restoration for improved cellular segmentation

**DOI:** 10.1101/2024.02.10.579780

**Authors:** Carsen Stringer, Marius Pachitariu

**Affiliations:** HHMI Janelia Research Campus, Ashburn, VA, USA

## Abstract

Generalist methods for cellular segmentation have good out-of-the-box performance on a variety of image types. However, existing methods struggle for images that are degraded by noise, blurred or undersampled, all of which are common in microscopy. We focused the development of Cellpose3 on addressing these cases, and here we demonstrate substantial out-of-the-box gains in segmentation and image quality for noisy, blurry or undersampled images. Unlike previous approaches, which train models to restore pixel values, we trained Cellpose3 to output images that are well-segmented by a generalist segmentation model, while maintaining perceptual similarity to the target images. Furthermore, we trained the restoration models on a large, varied collection of datasets, thus ensuring good generalization to user images. We provide these tools as “one-click” buttons inside the graphical interface of Cellpose as well as in the Cellpose API.

## Introduction

Generalist segmentation models like Cellpose do not work well out-of-the-box on degraded images, likely because there are very few noisy images in the training datasets. To perform segmentation on degraded images, one could first apply an image restoration method. Classical approaches to image restoration use various types of linear and nonlinear filtering [1–3], sometimes in combination with matrix decomposition methods [4–6]. Modern approaches instead use deep neural networks for various image restoration tasks, but this requires a training dataset of paired clean and noisy images because the networks are trained to predict clean images from noisy inputs [7, 8]. Neural networks for deblurring or upsampling typically work in the same way, although they sometimes require additional, image-specific knowledge such as the point-spread function [7–13].

The requirement for pairs of clean and corrupted images can be difficult to satisfy in practical experimental settings. To remove this requirement, methods like Noise2Self and Noise2Void learn to predict denoised pixel values from the context around each pixel [14, 15]. These methods do not require clean images, but rely on the independence of noise across pixels, and cannot be extended to other types of image restoration.

Another possible limitation of existing denoising techniques is their reliance on a pixel reconstruction loss, which may be suboptimal for subsequent image analysis tasks like segmentation. For example, minimizing the reconstruction loss can result in blurry images, similar to filtering approaches, because the network learns to return averages of possible cell shapes to reduce the pixel error [16]. In turn, blurry images may degrade segmentation quality, which relies on sharp outlines and shapes. Segmentation of noisy images has been used as an auxiliary task for denoising, but the corresponding cost functions were fully separated and trained with different neural networks [17].

In contrast to these previous approaches, here we asked whether images can be denoised or otherwise restored specifically *for* improved segmentation, and whether a single model trained on a varied dataset can generalize out-of-the-box to new images. We answer the first question by chaining together two neural networks: a trainable denoising network, followed by a pre-trained Cellpose segmentation network and we train the former network to minimize the cost function of the latter. In addition, we employ perceptual loss functions to make the reconstruction perceptually similar to the clean images [18, 19], and we use a balanced combination of large-scale datasets for training [20–26]. We start below by describing and demonstrating our approach in the denoising case; then we proceed to extend the approach to deblurring and upsampling.

## Results

### Model design and validation

Many imaging methods — especially fluorescence imaging — are affected by per-pixel noise which can substantially reduce the performance of segmentation algorithms, such as Cellpose (Figure 1a,b). This may be due to sensor noise, which is often described as Gaussian, or shot noise which is well captured by a Poisson distribution. Both types of noise can be avoided with higher stimulation intensities or longer exposures, but this often results in degradation of the sample or long imaging times which may be unacceptable. To create a diverse dataset of images with pixel noise, we added different amounts of Poisson noise to images of cells and other objects from the Cellpose test set [20]. Since Poisson noise becomes Gaussian-like in the limit of high simulated brightness, this approach should model both the sensor noise and the shot noise. To create a sensitive benchmark, we added sufficient noise to degrade the segmentation performance by a factor of *∼*2.

**Figure 1:**
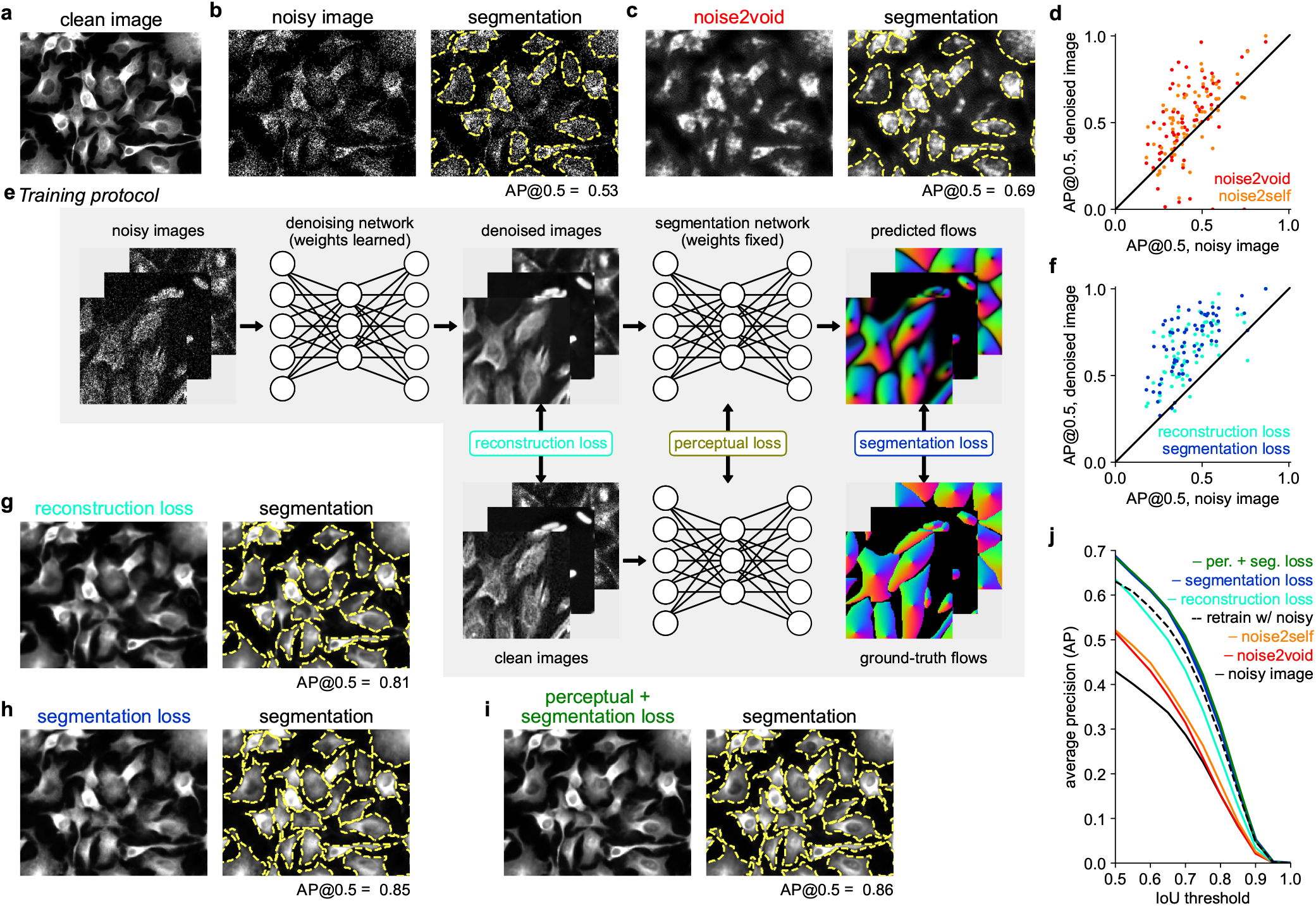
Generalist denoising model. **a**, Original (clean) image from the Cellpose test set. **b**, Same image with added Poisson noise (left) and segmentation (right). Average precision at an IoU threshold of 0.5 (AP@0.5) reported. **c**, Noisy image in **b** denoised with noise2self (left), and segmented (right). **d**, Improvement in segmentation performance after denoising with noise2void or noise2self for 68 test images. **e**, Training protocol for the Cellpose 3 denoising network. **f**, Same as **d**, for denoising with the reconstruction loss network (teal) or the segmentation loss network. **g-i**, Denoised images (left) and segmentations (right) for networks trained with different losses. **j**, Mean AP score as a function of IoU threshold across 68 test images, for the noisy and denoised images; dashed line shows a model directly trained to segment noisy images.

First, we applied two established techniques to the test data: Noise2Void and Noise2Self [14, 15]. These methods can blindly denoise images without requiring clean samples, which makes them directly applicable to our denoising test set, although they require training a new deep learning model in each case, which we did here (Figure 1c,d). When assessing the segmentation quality using the average precision (AP) metric (see Methods), we find that the denoised images indeed had higher segmentation accuracy compared to the noisy images, but the accuracy also decreased for a subset of the images (Figure 1d).

To improve on these approaches and remove the requirement of training at test time, our method learns a single denoising network for the entire Cellpose training set, which consists of diverse cellular and non-cellular images from various sources (Figure 1e, [27]). During training, Poisson noise of randomly-varied magnitude was added to the images on each training batch. We explored three different training objectives: the standard pixel reconstruction loss [7, 14, 15], a novel segmentation loss, and a perceptual loss [18, 19]. Training with the pixel reconstruction loss was effective at restoring image quality on a variety of test images, but sometimes removed or blurred the boundaries between cells, thus preventing successful segmentation (Figure 1g,j).

To improve on this, we reasoned that we can directly train the denoiser to output images that segment well, rather than images which reconstruct well (Figure 1e). We achieved this with a segmentation loss, computed by running the denoised images through the Cellpose ‘cyto2’ segmentation network [20] and comparing the outputs to the ground truth segmentations. The segmentation network remains fixed during training, and only the denoising network changes. This ensures that the denoising network outputs images that “look” right to the pretrained segmentation network. The training error thus backpropagates through the segmentation network, without changing it, and then further through the denoising network, producing gradients for learning. Training with the segmentation loss resulted in images where cell boundaries were more clearly distinguishable (Figure 1h,j). However, the restored images were still perceptually distinguishable from the clean images. While not detrimental for segmentation, the appearance may be distracting for human observers, especially when training new models with the human-in-the-loop approach from Cellpose2 [27].

To maintain visual fidelity to the ground truth images in a way that humans are sensitive to, we reasoned that a good approach would be to reconstruct abstract features of the images, rather than pixel values directly. This can be achieved with perceptual losses which aim to reconstruct the covariance matrix of feature activations in a pretrained neural network [18, 19]. Rather than using a standard ImageNet-pretrained model, we used the Cellpose model itself, which contains abstract information about cell appearance at different levels of its hierarchy (Figure 1e). Combining the perceptual and segmentation losses results in our final denoising network, which we will refer to as Cellpose3 (Figure 1i,j). As a further comparison, we retrained the original Cellpose model directly on the noisy images, to output segmentations without an intermediate denoising step. This approach did not perform as well as segmenting images that were denoised by Cellpose3 (Figure 1j).

The Cellpose3 denoising network successfully reconstructed a large variety of test images, not seen during training, recovering both the fine cell boundary details as well as the visual appearance of different types of microscopy data (Figure 2a, Figure S1a). One concern with all image reconstruction methods is that they could hallucinate cells on test images [28]. Our method allows for testing this possibility quantitatively by investigating whether false positive cells are detected on the reconstructed images. We found that the false positive rates were similar across all denoising approaches and similar to the false positive rate of the original noisy images (Figure S1b-d). Thus, we do not find quantitative evidence for hallucinations.

**Figure 2:**
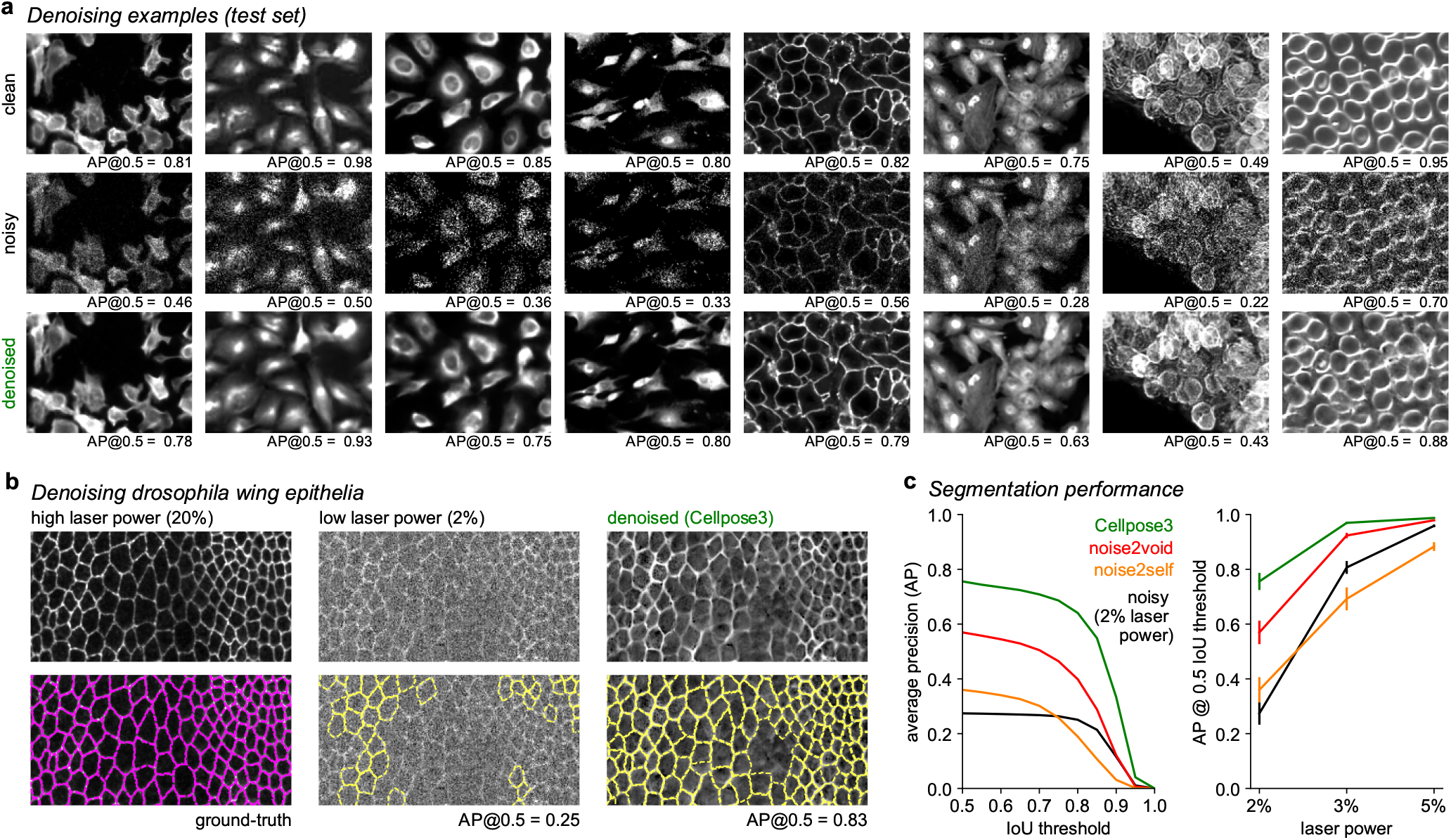
Denoising examples. **a**, Ground-truth images from the Cellpose test set (top), with Poisson noise added (middle), and denoised with Cellpose3. AP@0.5 reported. **b**, Top: Images of Drosophila wing epithelial cells [7] acquired with high laser power (left), low laser power (middle), and denoised with the Cellpose3 network (right). Bottom: segmentations. The high laser power segmentation was used as ground-truth. **c**, Left: mean AP score across 26 test images either original or denoised with various methods. Right: AP score at IoU threshold of 0.5 across laser powers. Error bars represent s.e.m., n=26 test images.

We also evaluated Cellpose3 on external data from [7], consisting of images of Drosophila wing epithelial cells acquired at different laser powers. First, we obtained segmentations of the images acquired at very high laser power using the original Cellpose1, which were near perfect by visual inspection (Figure 2b). We use these segmentations as ground truth for evaluating the reconstructions of images acquired at low laser powers. We find that Cellpose3 substantially increases the segmentation quality of these images, and more so than previous methods, across a range of laser powers (Figure 2b,c).

Finally, we wanted to ask how well Cellpose3 performs in cases where many images of the same type are available at test time. In such cases, retraining the blind denoising methods (Noise2Self, Noise2Void) may perform better compared to training on a single test image, which we confirmed on the Cell Image Library dataset CCDB:6843 consisting of 100 similar images [29] (Figure S2a). Nonetheless, the Cellpose3 approach still outperformed these previous approaches both in visual quality and segmentation quality (Figure S2b,c). In addition, we trained CARE using pairs of noisy and clean training images from the training data of the specialized dataset [7]. Cellpose3 also outperformed this approach.

We next applied this approach to a large “Nuclei” dataset of nuclear images from various sources [23, 30–32], which were previously used to train the Cellpose ‘nuclei’ segmentation model [20] (Figure 3a, Figure S3a). In this case, we found that all three losses (reconstruction, perceptual, segmentation) produced almost identical segmentation performance on the test set, which again outperformed the blind denoising methods (Figure 3b). We suspect this is because nuclear segmentation relies on simpler object shapes which may be easier to denoise. Similar to the cellular denoisers, the nuclear denoisers did not hallucinate nuclei, as quantified by the false positive, true positive and false negative rates (Figure S3b-d). On images of Tribolium nuclei acquired at different laser powers from [7], the Cellpose3 denoised images produced segmentations which matched the ground-truth segmentations significantly better than the other denoising approaches (Figure 3c,d).

**Figure 3:**
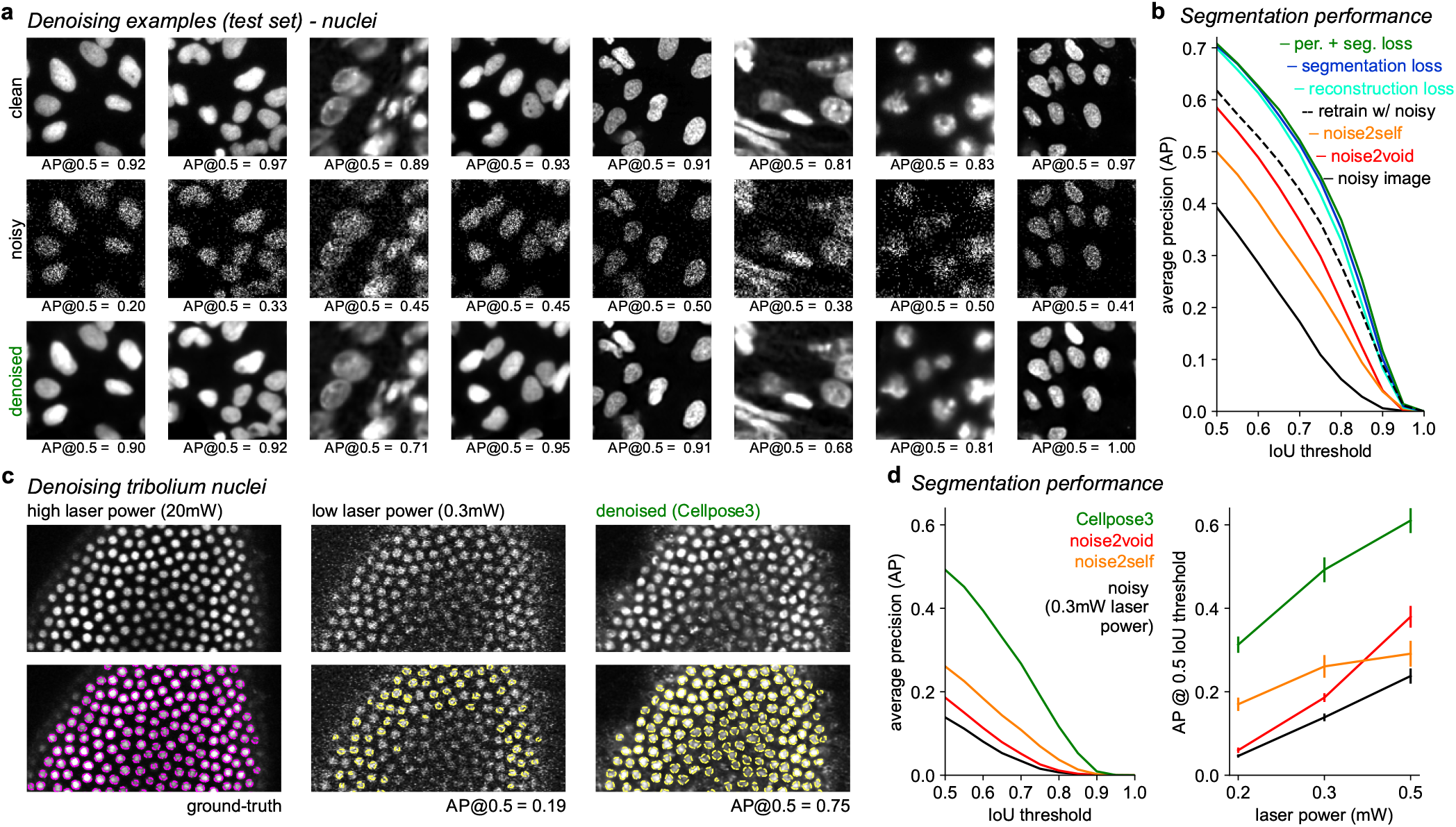
Denoising nuclei. **a**, Example ground-truth image crops from the Nuclei test set (top), with Poisson noise added (middle), and denolised (bottom). AP@0.5 reported for entire, uncropped image. **b**, Mean AP score averaged across 111 test images for noisy and denoised images; dashed line shows a model directly trained to segment noisy images. **c**, Images of Tribolium nuclei during development [7], acquired with high laser power (left), low laser power (middle), and denoised with Cellpose 3 (right). Bottom: segmentations. The high laser power segmentation was used as ground-truth. **d**, Left: mean AP score across 6 test images either original or denoised with various methods. Right: Mean AP score at IoU threshold of 0.5 across laser powers. Error bars represent s.e.m., n=6 test images.

### Deblurring and upsampling

Next we considered whether the same approach can be applied to other image restoration tasks, such as deblurring and upsampling. Blurring in microscopy is typically due to scattering such as in thick tissue imaging. Computational deblurring methods — sometimes referred to as “deconvolution”— can reveal finer details than are visible in the blurred images. Similarly, under-sampling of a field of view is also common, often due to speed or photon-budget constraints. Upsampling such images —sometimes referred to as “super-resolution”— can restore details that make the segmentation better.

For deblurring, we trained our networks on images with Gaussian blur of random spatial sizes. For test images, we again fine-tuned the amount of degradation per image to approximately decrease the segmentation performance by half. Networks trained on the Cellpose training set were able to deblur a variety of cellular images at test time (Figure 4a), improving the segmentation performance substantially (Figure 4b). We also trained the networks on the Nuclei training set and found a large improvement in segmentation performance after deblurring (Figure S4a,b). The networks trained with both the perceptual and segmentation loss again performed best (Figure 4b, Figure S4b).

**Figure 4:**
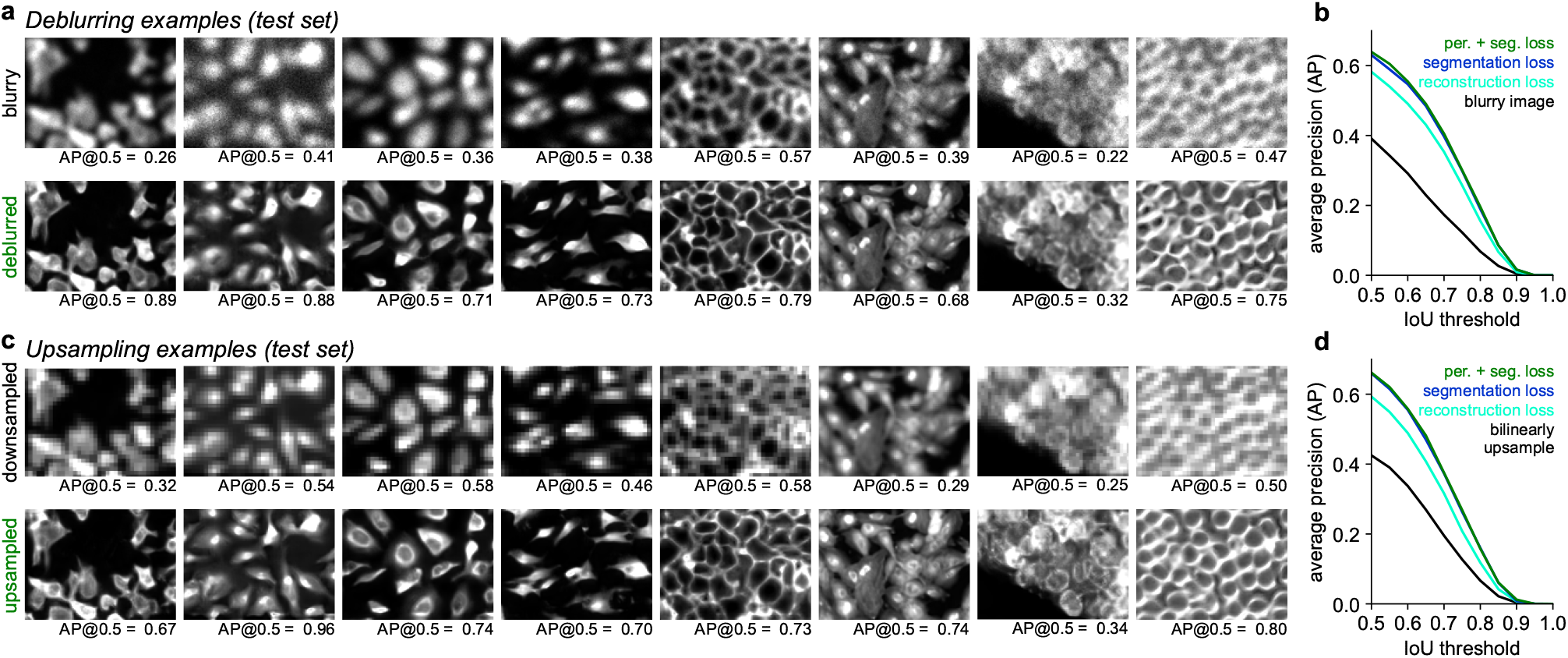
Deblurring and upsampling cellular images. **a**, Same clean image crops as shown in Figure 2a, with Gaussian blurring (top), and deblurred with Cellpose3 (bottom). AP@0.5 is reported for entire image. **b**, Mean AP score across 68 test images either blurry or deblurred. **cd**, Same as **ab** for downsampled images that were upsampled with a Cellpose3 network.

For upsampling, we trained networks on images that were downsampled by decimation. The test set was again constructed to approximately reduce segmentation performance in half. The upsampling networks trained on the Cellpose dataset recovered some of the visual detail in the test images lost through downsampling and improved segmentation (Figure 4c,d). The networks trained on the Nuclei dataset also produced upsampled images with improved segmentation performance (Figure S4c,d). As before, networks trained with both the perceptual and segmentation loss performed best.

### One-click image restoration

So far we have shown separate models trained on cells and nuclei. There are other datasets that we could have used such as TissueNet [22], LiveCell [21], Omnipose [24], Yeaz [26], DeepBacs [25]. Instead of training new restoration models in each case, we asked whether single restoration models could be trained on all these datasets combined. Note, however, that we still train individual models for each of the three restoration tasks; combining tasks into a single model was in fact detrimental to performance (results not shown).

Training a single restoration model across all datasets required first training a super-generalist “cyto3” segmentation network, to be used for computing the segmentation and perceptual losses (Figure S5). Similar to the Cellpose2 paper [27], we observed a small but consistent loss in performance from the network trained on all datasets (for example, AP@0.5 on the Cellpose test set was 0.786 versus 0.771 for the dataset-specific versus the super-generalist ‘cyto3’ model). This result differs markedly from a recent study [33], which found a much larger decrease in performance for generalist versus specialist Cellpose models, likely due to the suboptimal training regimen used there (see Methods for more details). Here we adjusted the sampling probability of the datasets during training, to prevent large but homogeneous datasets like TissueNet and LiveCell from dominating the statistics of the cost function. We also tested a transformer backbone for Cellpose and found that it did not improve performance [34, 35]; in fact, it had nearly identical performance to our residual U-net backbone, which suggests that generalization performance is limited by dataset diversity rather than model architecture.

We trained separate networks for denoising, deblurring and upsampling of all the training images from the combined 9 datasets. This resulted in networks which performed as well as the dataset-specific models in all cases except for deblurring and upsampling of nuclei (Figure 5a,b). The models also worked well on the other 7 datasets (Figure 5c). We refer to these as “one-click” models, as they can be run simply by pressing the corresponding button in the Cellpose GUI without specifiying the image type. We also provide the dataset-specific nucleus model which should be useful especially for deblurring and upsampling.

**Figure 5:**
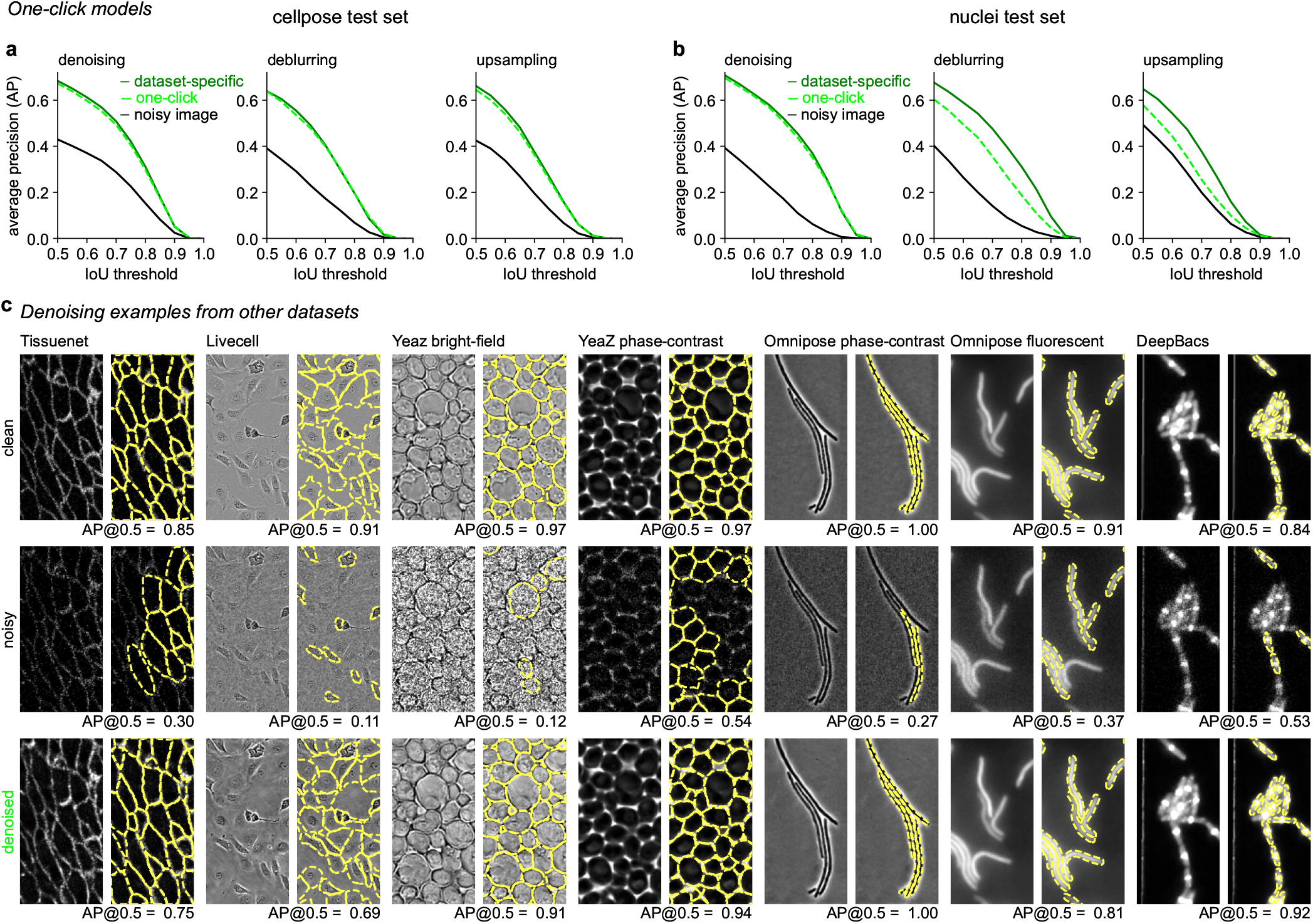
One-click models for denoising, deblurring and upsampling. **a**, Segmentation performance on the Cellpose test set images after denoising (left), deblurring (middle), or upsampling (right) with the one-click models compared to the dataset-specific models. **b**, Same as **a** for the Nuclei test set. **c**, Example images and their segmentations from other datasets (top), with Poisson noise added (middle) and denoised with the one-click model (bottom). AP@0.5 reported.

## Discussion

Here we introduced a new method for denoising biological images, which focuses on improving a downstream task such as segmentation, rather than image reconstruction. Like the blind deconvolution methods [14, 15], our approach does not require clean images at test time. By training this method on a large and diverse dataset, we achieved high out-of-sample generalization on new images, which should enable users to easily employ the Cellpose3 denoisers on their own data.

Although our method focuses on segmentation rather than reconstruction, the restored images are nonetheless visually compelling and can be used to create manual annotations with the human-in-the-loop approach from Cellpose2. Note that we do not report here some of the standard denoising metrics like PSNR, since these metrics are essentially a different way to estimate the per-pixel reconstruction loss. Our approach is not designed to reproduce exact pixel values, which may in general be impossible from limited information, but rather to reproduce more abstract features of the cells, such as their boundaries and their perceptual appearance.

It is also important to discuss the use of denoised images in scientific research. Since modern denoisers use complex image priors learned from their training datasets, there is always a risk that certain image features are “invented” by the network rather than truly being present in the data. With respect to Cellpose3, one could take two approaches: 1) a conservative approach, where Cellpose3 is used to detect and segment cells, but the biological quantifications are done on the raw images; 2) a liberal approach, where all analyses are done on the denoised images. We believe the conservative approach should be used whenever possible. When this is not possible, the liberal approach could be used if well-matched control images are available to verify the method. For example, pairs of high/low quality images could be obtained for a small subset of the data to serve as an internal control.

## Acknowledgments

This research was funded by the Howard Hughes Medical Institute at the Janelia Research Campus. Thanks to Michael Rariden for contributions to the Cellpose code base.

## Author contributions

C.S. and M.P. designed the study, performed data analysis, and wrote the manuscript.

## Data availability

No new data was generated in this study. The ‘cyto2’ dataset is publicly available at https://www.cellpose.org/dataset, and the other datasets were generated and shared by other labs [21–26, 30–32].

## Code availability

Cellpose3 was used to perform all analyses in the paper. The code and GUI are available at https://www.github.com/mouseland/cellpose. Scripts for recreating the analyses in the figures are available at https://github.com/MouseLand/cellpose/tree/main/paper/3.0.

## Methods

The Cellpose code library is implemented in Python 3 [36], using pytorch, numpy, scipy, numba, opencv, imagecodecs, tifffile, fastremap, and tqdm [37–45]. The graphical user interface additionally uses PyQt, pyqtgraph, and superqt [46–48]. The figures were made using matplotlib and jupyter-notebook [49, 50].

### Cellpose3 denoising network training and testing

#### Model architecture

We used the same model architecture as the Cellpose model, described in detail in [20]. The Cellpose model is a deep neural network with a U-net based architecture, consisting of four downsampling blocks and four upsampling blocks, each block containing four convolutional layers with residual connections [51, 52]. The model takes as input a single noisy image and outputs the image denoised. On 224×224 images, the network runs at a speed of 750 images per second on a server A100 GPU, or 210 images per second on a consumer-grade A4000 GPU.

#### Training

All training was performed with the AdamW optimizer [53]. The learning rate increased linearly from 0 to 0.001 over the first 10 epochs, then decreased by factors of 2 every 10 epochs over the last 100 epochs. Each network was trained for 2000 epochs. The one-click models were initialized using the weights from the Cellpose3 networks trained with the ‘cyto2’ dataset; all other models were trained from scratch.

The reconstruction loss was the mean squared error between the ground-truth image and the predicted image (the network output). To compute the segmentation loss, we used the ‘cyto2’ model, the ‘nuclei’ model, or the new super-generalist model. The segmentation loss was the same loss as the Cellpose segmentation loss [20]: the mean squared error between the XY flows from the ground-truth segmentation and the predicted XY flows, scaled by a factor of five, added to the binary cross-entropy between the ground-truth cell probability and the the predicted cell probability. For the perceptual loss we computed the correlation matrix of neural network activations at each downsampling block [18, 19]. A target correlation matrix was computed for the ground truth image, and a predicted correlation matrix was computed for the denoised/deblurred/upsampled image. The perceptual loss function was defined as the mean squared error between these two matrices, normalized by the standard deviation of the target correlation matrix in each downsampling block. This latter normalization ensures that each block level contributes approximately the same variance to the cost function.

All images were normalized such that 0 was set to the first percentile of the image intensity and 1 was the 99th percentile of the image intensity. The image intensities were then clipped at 0 – this was necessary to add Poisson noise.

During training, all images were resized such that the cell or nuclei diameters were 30.0 or 17.0 pixels respectively. Then, before adding synthetic noise, the images were resized between 0.5 to 2 times their original size, and randomly cropped to a size of 340×340. This allowed the Poisson noise to be added at various scales. After adding the noise, blurring and/or downsampling, the images were randomly resized and cropped to 224×224 using the original Cellpose augmentations: random rotation, random flipping, and random resize with a scale factor between 0.75 and 1.25, such that 1.0 is equivalent to cell or nuclei diameters of 30.0 or 17.0 pixels. When training the one-click models using the super-generalist segmentation model, all images were resized such that the cells or nuclei diameters were 30.0 pixels. When training the one-click models, the images across all datasets were sampled during each epoch, as described in the super-generalist training section.

#### Synthetic noise

We added three types of perturbations to images: Poisson noise, Gaussian blurring, and downsampling. During training, for the Poisson noise generation, we multiplied each image by a random scaling factor, drawn from a gamma distribution with α=4.0 and varying values of β, and then used this scaled image to draw a random sample of Poisson noise. Poisson noise was generated on 80% of the training images, with β=0.7 during denoising training, β=0.1 during deblurring training, and β=0.01 during upsampling training. This operation was applied last.

For the deblurring and upsampling training, we blurred the images 80% of the time with an isotropic 2D Gaussian filter with a standard deviation drawn from an exponential distribution with a mean of 0.5, clipped between 0.1 and 1.0. These standard deviations were then scaled proportional to the diameter of the cells/nuclei in the training image, with a scaling factor of 1/3 for deblurring and 1/6 for upsampling, i.e. for images scaled to an average diameter of 30.0, the average blurring scaling factor would be 10.0 for deblurring and 5.0 for upsampling. This operation was applied first.

For the upsampling training, we downsampled the images 80% of the time, with a downsampling factor drawn as a random integer between 2 and 7. The images were downsampled using subsampling with the random integer, then bilinearly interpolated to the original size to be input into the network. This operation was applied after blurring and before Poisson noise was added.

For testing, we added different amounts of noise/blurring/downsampling to each image, scaled to reduce the segmentation performance of ‘cyto2’ or ‘nuclei‘ on the image by 50%, as measured by the average precision at an IoU threshold of 0.5. The scaling factors for Poisson noise varied from 0.5 to 80.0, the Gaussian blurring standard deviations varied from 0.5 to 10.0, and the downsampling factors varied from 1 to 6. When generating the blurry images, a small amount of Poisson noise was added, with a scale factor of 120.0. When generating the downsampled images, blurring was performed with a standard deviation proportional to 1/2 the downsampling factor. After these operations, each of the noisy images was normalized such that 0 was the first percentile and 1 was the 99th percentile.

#### Images with real noise

We used two imaging datasets from [7] collected at varying laser powers to demonstrate the ability of Cellpose3 to denoise images with real Poisson shot noise (Figure 2b,c, Figure 3c,d). The Drosophila wing epithelia cell dataset consisted of 26 images collected at four different laser powers. Each image consisted of multiple planes – we used the maximum projection image provided in the paper. The Tribolium nuclei dataset consisted of 6 images collected at four different laser powers. Each image consisted of multiple planes – we used the maximum projection image across 16 planes in the middle of the volume. Ground-truth segmentations for the images were not provided, so we used the Cellpose segmentations of the high laser power images as the ground-truth. Based on the average ROI size in the high laser power segmentation, we set the diameters to 24 and 12 for the epithelia and nuclei datasets respectively, and used these values for all laser powers and for the denoised images. For the Drosophila wing epithelia cells (Figure 2b,c), we set the cell probability threshold to -2.0 for all segmentation evaluations to ensure the masks filled the cytoplasmic space across noise levels and the denoised images.

### Segmentation networks

#### Cellpose model

The Cellpose model architecture is described in detail in [20]. The Cellpose model predicts 3 outputs: the probability of a pixel being inside a cell (1), the flows of pixels towards the center of a cell in X (2) and Y (3). The flows are then used to construct the cell masks. The Cellpose model, ‘cyto2’, was trained on 796 images of cells and objects with 1 or 2 channels (if the image had a nuclear channel), with an average cell diameter of 30.0 pixels used in training. The Cellpose model, ‘nuclei’, was trained on 1025 single-channel images of nuclei, with an average nucleus diameter of 17.0 pixels used in training.

#### Noisy image training

The ‘retrain w/ noisy’ networks were Cellpose networks trained from scratch in the same way as the Cellpose 1.0 paper: 500 training epochs with a batch size of 8, stochastic gradient descent optimizer, weight decay of 0.00001, momentum of 0.9, and learning rate of 0.2. The learning rate increased linearly from 0 to 0.2 over the first 10 epochs, then decreased by factors of 2 every 10 epochs after the 400th epoch. The proportion and scaling of noise added to the images was the same as what was added to the Cellpose3 networks trained on the same noise type.

#### Dataset-specific, super-generalist, and transformer training

The dataset-specific and super-generalist models use the same architecture as the original Cellpose model which we found to be sufficient. Increasing the width of the layers or the depth of the network did not help (results not shown). Recent analyses suggest that super-generalist models *require* transformer models and perform better than Cellpose [33, 54], but we believe the comparisons have not been done correctly. For example, [54] allows the transformer model to retrain but does not retrain Cellpose. As another example, [33] and [55] trained Cellpose using the finetuning protocol from the Cellpose2 paper, rather than the protocol for training Cellpose from scratch, and the training dataset was dominated by TissueNet images which resulted in class imbalance, a known problem for training supervised models. It may be that transformers are less affected by such training set imbalance, but more likely the very long training times of transformers compensate for the class imbalance. Even transformers can benefit from class balancing, as the winner of the NeurIPS 2022 cell segmentation challenge (MEDIAR [34]) used a complex balancing strategy.

To make the comparison to transformers more direct, we replaced only the backbone of Cellpose with a transformer, keeping everything else the same. This is similar to the approach of the MEDIAR algorithm. This transformer consisted of a transformer-based segformer encoder and a multi-scale attention decoder, which was trained to predict the cell flows and probabilities from Cellpose [56, 57]. As in the MEDIAR paper, we used the implementation of the encoder and decoder from the segmentation models.pytorch github [35]. Also, as in the MEDIAR paper, we used the MIT-B5 segformer encoder and initialized the encoder with weights from pretraining on imagenet provided in the github repository.

To optimize these models, we used the AdamW optimizer [53]. The learning rate increased linearly from 0 to 0.005 over the first 10 epochs, then decreased by factors of 2 every 10 epochs over the last 100 epochs. The super-generalist and transformer models were trained for 5000 epochs, and the dataset-specific model was trained for 2000 epochs. For the super-generalist and transformer models, each epoch consisted of 800 training images, randomly sampled from the total training set size of 8402 images. For the dataset-specific models, each epoch consisted of all training images in the dataset, if there were fewer than 800 training images, or consisted of 800 images randomly sampled from the dataset. Each image had one or two channels, where the second channel contained the nuclear channel if it was a cellular image with two channels, or otherwise was set to zero. The nuclei-only images had the nuclear channel as the first channel and the second channel set to zero. We used the original Cellpose augmentations, as described in the denoising section, with an average cell/nuclei diameter of 30.0 pixels for all images.

For the super-generalist and transformer models, the sampling probability of each image varied depending on the image set: PhC yeast images and fluorescent bacterial images were sampled at a probability of 1% each; bright-field yeast images, phase bacterial images, and DeepBacs images at 2% each; livecell images at 5%; tissuenet images at 8%; nuclei images at 20%; and cyto2 images at 59%. We upweighted images in the cyto2 and nuclei training sets because they contained the most variability across images. These sampling probabilities and 800 images per epochs were also used when training the super-generalist size model and also for the one-click model training. For the size network training, we used the same defaults as in the Cellpose paper: 10 epochs of training images and an L2 regularization constant of 1.0 [20].

The flow representations of the masks were modified slightly from the original method [20]. We used the point within the mask closest to the center-of-mass as the source of the diffusion process, instead of the median. In cells with a long process, like a dendrite, for example, this puts the source closer to the process rather than in the soma, to improve convergence of such longer processes. We also no longer logarithmically scaled the diffusion probabilities before computing the gradients, and when normalizing the gradients by their L2 norm, we added a smaller epsilon to the L2 norm (1e-60 vs 1e-20). This increased the size of smaller gradients, improving the performance on long and thin cells.

#### Evaluation

The Cellpose segmentation models are trained such that all cells and nuclei are approximately the same size in pixels across all images, by resizing each image such that the average cell (or nuclei) diameter is 30.0 (or 17.0). Therefore, the images in the test set must be resized before running the segmentation models. In all analyses besides Figure S5, the test images were rescaled per image using the average diameter computed from the ground-truth masks, rather than using the size estimation models from Cellpose. This was done because the size estimation models were trained on images that were not noisy, blurry or downsampled, and so would not perform correctly on those images, and also because the objective of the analysis is to assess the segmentation – we assume most users know the diameters of their cells or nuclei in their images and can provide them for the denoising and segmentation algorithms.

In Figure S5, we evaluated the segmentation performance of the super-generalist model versus other models. For this, we did not assume that the network knows the average diameter of the cells/nuclei in the image and therefore used the super-generalist model to create a size model to estimate the diameters. Using these estimated diameters, each image was resized to set the average diameter to approximately 30 pixels, and then run through each of the different segmentation models.

The flow error threshold (quality control step) was set to 0.4 and the cell probability threshold was set to 0. In Figure S5, for all models we turned on test-time augmentations during evaluation, as in [20] and [34]. When segmenting the phase-contrast images from the Omnipose dataset, we set the number of iterations ‘niter’ for the dynamics post-processing to 2000 for all images, to improve the convergence for long and thin cells.

#### Quantification of segmentation quality

As described in Cellpose 1 and 2, we quantified the predictions of the segmentation algorithms by matching each predicted mask to the ground-truth mask that is most similar, as defined by the intersection over union metric (IoU) between the predicted and ground-truth. The average precision metric (*AP*) for each image is defined using the true positives (matches with IoU above a given threshold), false positives (predicted masks without matches), and false negatives (missed ground-truth masks):

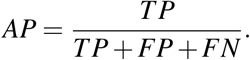

The average precision reported is averaged over the *AP* for each image in the test set. We computed the true positive rate, false positive rate and the false negative rate by normalizing the TP, FP and FN values per image by the total number of ground-truth masks per image, then averaging across images.

### Denoising model comparisons

We compared the performance of the Cellpose3 models to Noise2Void, Noise2Self, and CARE [7, 14, 15]. In Figure 1 and Figure 3a,b, we trained Noise2Void and Noise2Self on a per image basis, as described in their papers, creating 68 models for the 68 noisy cellular test images and 111 models for the 111 noisy nuclear test images. In Figure 2b,c and Figure 3c,d, we also trained Noise2Void and Noise2Self on a per image basis on each of the three noise levels, creating 78 models for the 78 noisy Drosophila epithelia test images and 18 models for the 18 noisy Tribolium nuclei test images. Each of these images was normalized such that 0 was the first percentile and 1 was the 99th percentile.

For specialist training, in Figure S2, we trained and tested Noise2Void, Noise2Self, and CARE on images from the CellImageLibrary dataset CCDB:6843 [29], consisting of 89 images in the training set and 11 images in the test set. We added 20 instances of random noise to each training image, drawn from the same distribution as used in Cellpose3 training, and normalized in the same way. We then created a validation set from these images using quarter crops: 1/4 of each image was used for validation and the other 3/4s for training, resulting in 3 training crops per image. In total this resulted in 5340 noisy training images and 1780 noisy validation images. Each training and validation image had a ground-truth pair – the original image without noise – which was used for training CARE. Noise2Void and Noise2Self were additionally trained on the test images.

#### Noise2Self model

Noise2Self is a blind denoising algorithm which does not use clean images for training, and can be trained on single images [14]. For the single noisy image training, we first tried the ‘Single-Shot Denoising’ notebook in the Noise2Self github, using the default CNN model defined with 8 layers and the default training parameters: Adam optimizer, learning rate of 0.01, and 500 epochs. At test time, we used the model’s output from the epoch with the lowest validation loss, which is defined as the mean squared error between the masked pixels from the noisy image and the masked pixels from the model output (as is done in the notebook). However, we found that the average segmentation performance on the denoised images from this training was poor (average AP@0.5 of 0.435 for cellular images). Instead, we used the U-net model used for CellNet training, as described in the Noise2Self paper. We used a batch size of 8, and the same augmentations as Cellpose: random rotation, cropping and resizing (scale between 0.75 to 1.25), and flipping, with random crops of size 128 by 128. We trained each single image model with the Adam optimizer with a learning rate of 0.0005 and for 100 epochs. At test time, the test images were padded with zeros to achieve image dimensions that were divisible by 16 so that they could be run through the U-net, and then the padding was cropped out on the output. This resulted in an average AP@0.5 of 0.511.

For the specialist training, we used the U-net model and a batch size of 64, as used for the CellNet training. We used the same transformations as the per image training. We performed a sweep over learning rates (0.00001 - 0.01) to determine the best learning rate on the validation set based on the validation loss (mean-squared error). We found a learning rate of 0.0005 produced the lowest validation loss. We also tried different numbers of epochs, but found that, although training for more epochs reduced the validation loss, the test set segmentation performance became worse with more epochs. Therefore, we trained for 50 epochs, the setting used for the CellNet training. As in the per image training, the images were padded then cropped during test time to enable them to be run through the U-net.

#### Noise2Void model

Noise2Void is also a blind denoising algorithm which does not use clean images for training, and can be trained on single images [15]. For the per image training we used the parameters and model from the denoising2D SEM training notebook. This default model was a U-net with a depth of 2 and a kernel size of 3. The training used a validation set, so we divided each noisy test image into quarters and used one quarter for validation. As suggested in the notebook, we set the number of epochs to 100, train steps per epoch of 1, learning rate of 0.0004, batch size of 128 (or less depending on the number of patches), and a patch size to 64×64 (for some images it was required to be size 60×60). We used the default augmentations, as defined in the function ’generate patches from list‘: flipping and rotating by 90 degrees, and all possible crops of the image into 64×64 patches.

For the specialist training, we used the same U-net model, the same augmentations and the default batch size of 128. We used the same transformations as the per image training and used 25 training steps per epoch. We performed a sweep over learning rates (0.00001 - 0.01) to determine the best learning rate on the validation set based on the validation loss (mean-squared error). We found a learning rate of 0.0004 (default) produced the lowest validation loss. As with Noise2Self, we also tried different numbers of epochs, but found that, although training for more epochs reduced the validation loss, the test set segmentation performance became worse with more epochs. Therefore, we trained for 100 epochs, the setting recommended in the notebook.

#### CARE model

The CARE model is a denoising algorithm trained to restore noisy images using noisy-clean image pairs. We only performed specialist training for CARE as it cannot be trained per image on noisy images.

For specialist training, we used the default CARE U-net model with a depth of 2 and a kernel size of 3, provided in the denoising2D training example notebook. We used the default parameters: two 128×128 patches per training/validation image, a training batch size of 8, and the number of training steps per epoch of 400 (equivalent to the number of images per epoch as Noise2Void). We turned off image normalization in CARE, as it was similar to our normalization, and when on, it resulted in a subset of images with pixel values over 1e20. We performed a sweep over learning rates (1e-5 - 1e-2) to determine the optimal hyperparameters on the validation set based on the validation loss (mean-squared error). We found a learning rate of 1e-3 and 100 epochs produced the lowest validation loss, and using these hyperparameters chose the network with the lowest validation loss across training epochs (as is default in the CARE code). We then ran the test images through the network’s ‘predict’ function.

### Datasets

We used 9 publicly available datasets for training the one-click restoration models, and for training the super-generalist ‘cyto3’ model.

#### Cellpose cyto2 dataset

This dataset consists of the 618 images from the ‘cyto’ dataset (540 training images and 68 testing images), described in detail in [20], from various sources [29, 58–61], and an additional 256 new training images. The new training images were found using Google image searches or were submitted by users of Cellpose. The dataset is available at https://www.cellpose.org/dataset.

#### Cellpose nucleus dataset

This dataset of nuclear images was described in detail in [20], it consists of 1025 training images and 111 test images from various sources [23, 30–32].

#### TissueNet

The TissueNet dataset consists of 2601 training and 1249 test images of 6 different tissue types collected using fluorescent microscopy on 6 different platforms (https://datasets.deepcell.org/) [22]. This dataset includes nuclear and cellular segmentations for each image – we used the cellular segmentations.

#### LiveCell

The LiveCell dataset consists of 3188 training and 1516 test images of 8 different cell lines collected using phase-contrast microscopy (https://sartorius-research.github.io/LIVECell/) [21]. The images were segmented with overlaps allowed across masks. The Cellpose model cannot predict overlapping masks, so overlaps were removed, as described in the Cellpose 2.0 paper [27].

#### Omnipose

The Omnipose dataset includes of two bacterial datasets, each with manual segmentations: fluorescent bacterial images (143 training and 75 test images), and phase-contrast microscopy bacterial images (249 training and 148 test images) [24].

#### YeaZ

The YeaZ dataset consists of two datasets of manually-segmented yeast cells: phase-contrast images (16 2D training images and 6 2D test images), and bright-field images (229 training images and 77 test images) [26].

#### DeepBacs

The DeepBacs dataset consists of 155 training images and 35 test images of bacteria collected using bright-field microscopy and fluorescent imaging [25]. We excluded 5 test images when computing segmentation quality, because they were labeled in a different modality – this only had a small effect on the AP score.

---

**Figure S1:**
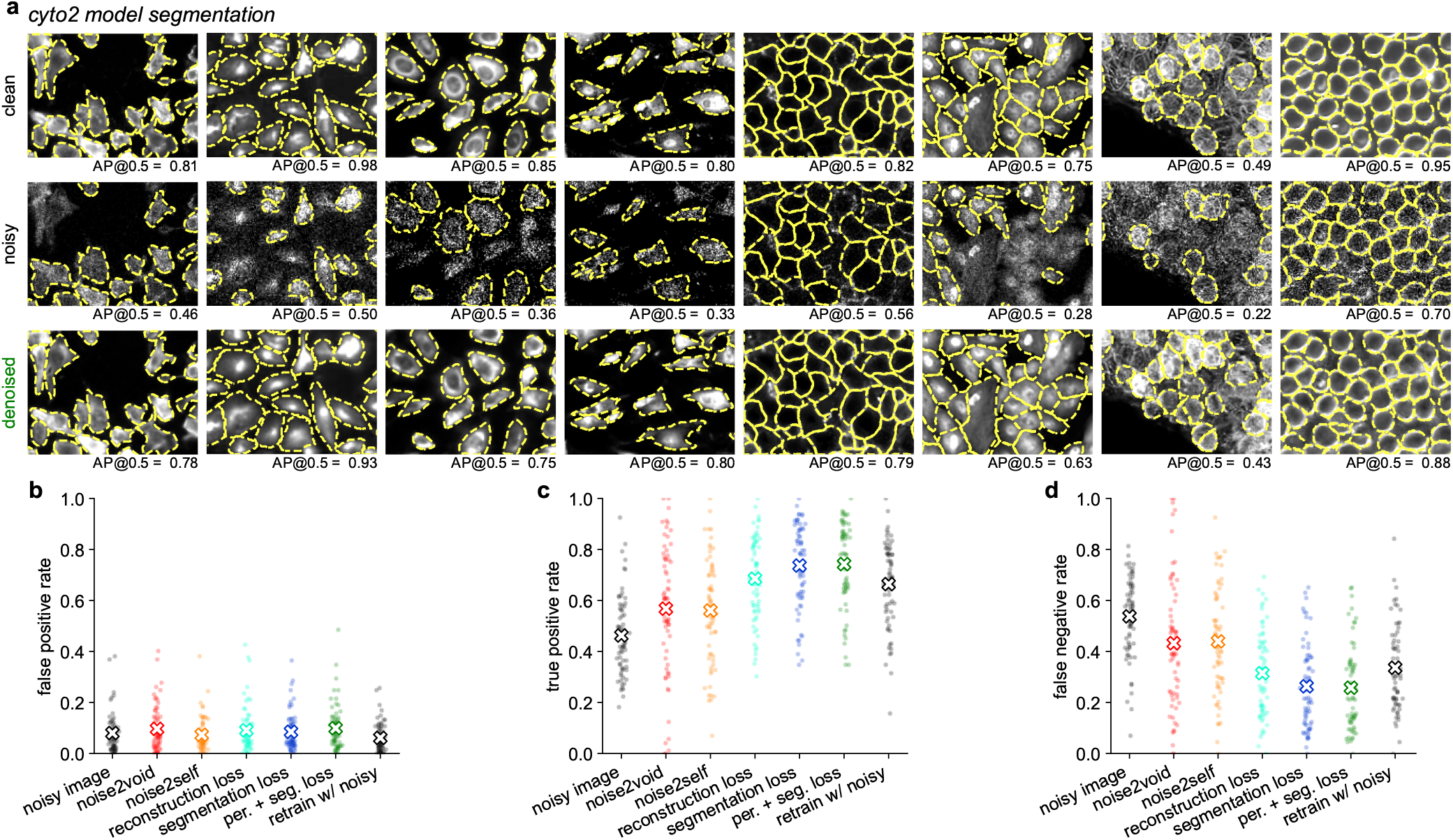
Segmentation of clean, noisy and denoised images of cells. **a**, Same as Figure 2a but overlaid with segmentations. **b**, The false positive rate of the segmentation at an IoU threshold of 0.5 for each of the 68 test images, either denoised and segmented with ‘cyto2’ or segmented directly with a model retrained on noisy images. Averages marked with ”x”. **cd**, Same as **b** for the true positive rates and the false negative rates respectively.

**Figure S2:**
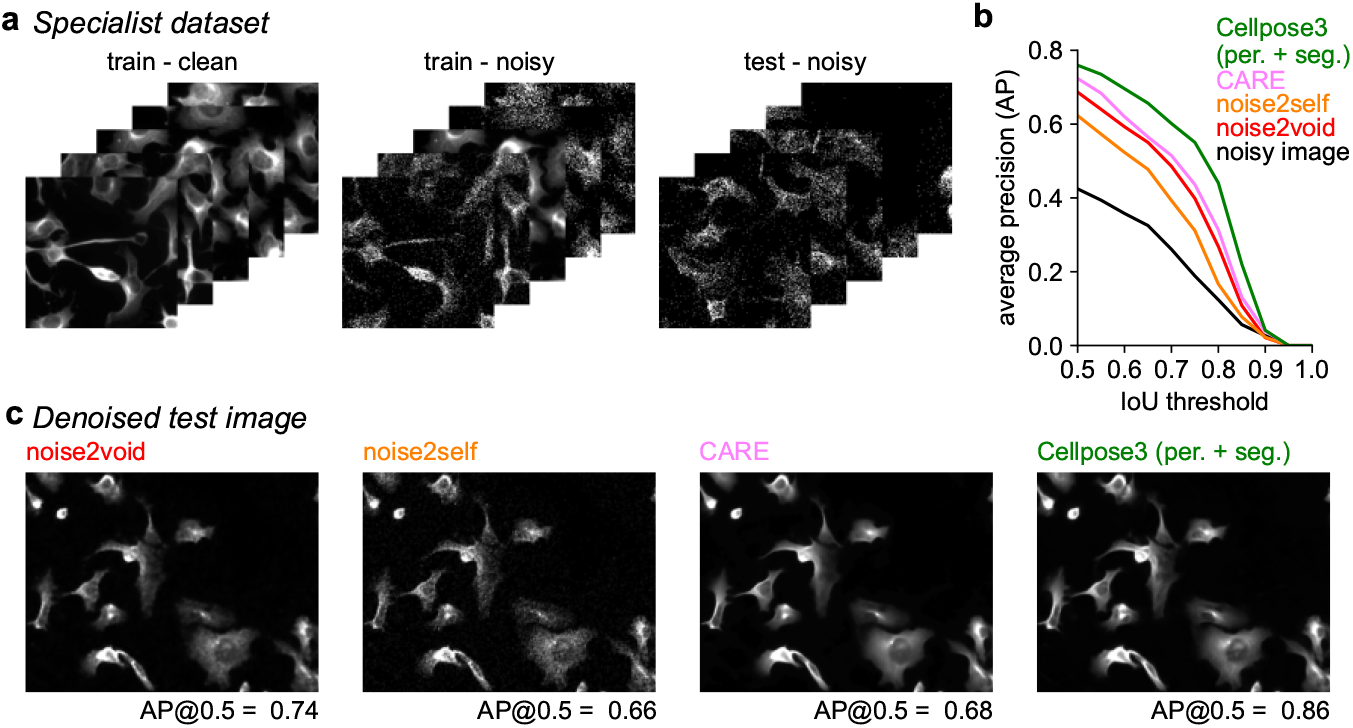
Denoising a specialist dataset. **a**, 89 training images, clean and with Poisson noise added, and 11 test images with Poisson noise, from the CellImageLibrary CCDB:6843 dataset [29]. **b**, Mean AP score of segmentation performed using the ‘cyto2’ model applied to the noisy and denoised images, averaged across 11 test images. The CARE model was trained using the 89 noisy and ground-truth image pairs, and Noise2Self and Noise2Void were trained on the 100 noisy images from both the training and test set. The Cellpose3 model was the same as the model in Figure 1, which was trained using all ‘cyto2’ images.

**Figure S3:**
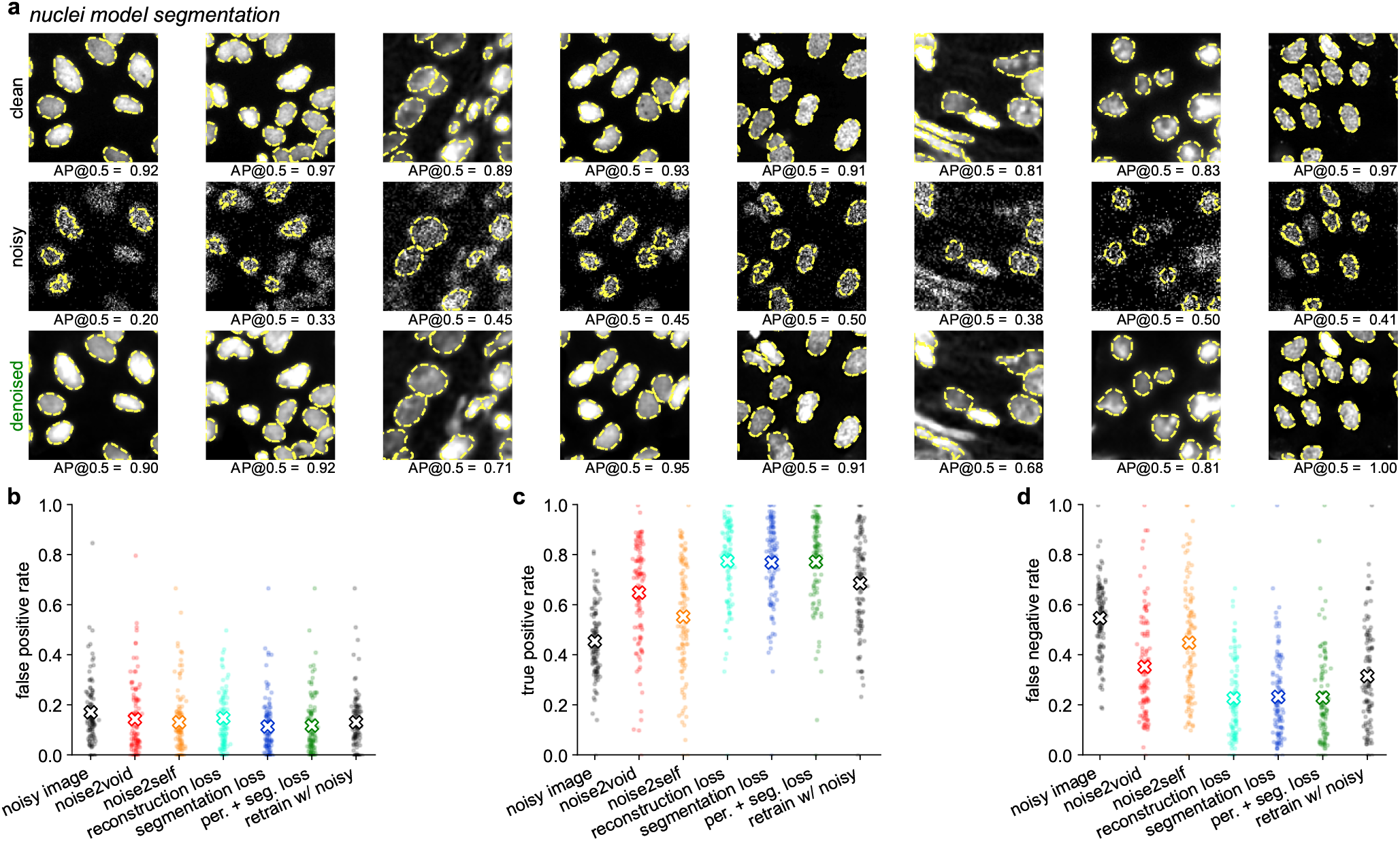
Segmentation of clean, noisy and denoised images of nuclei. Same as Figure S1, for the Nuclei dataset.

**Figure S4:**
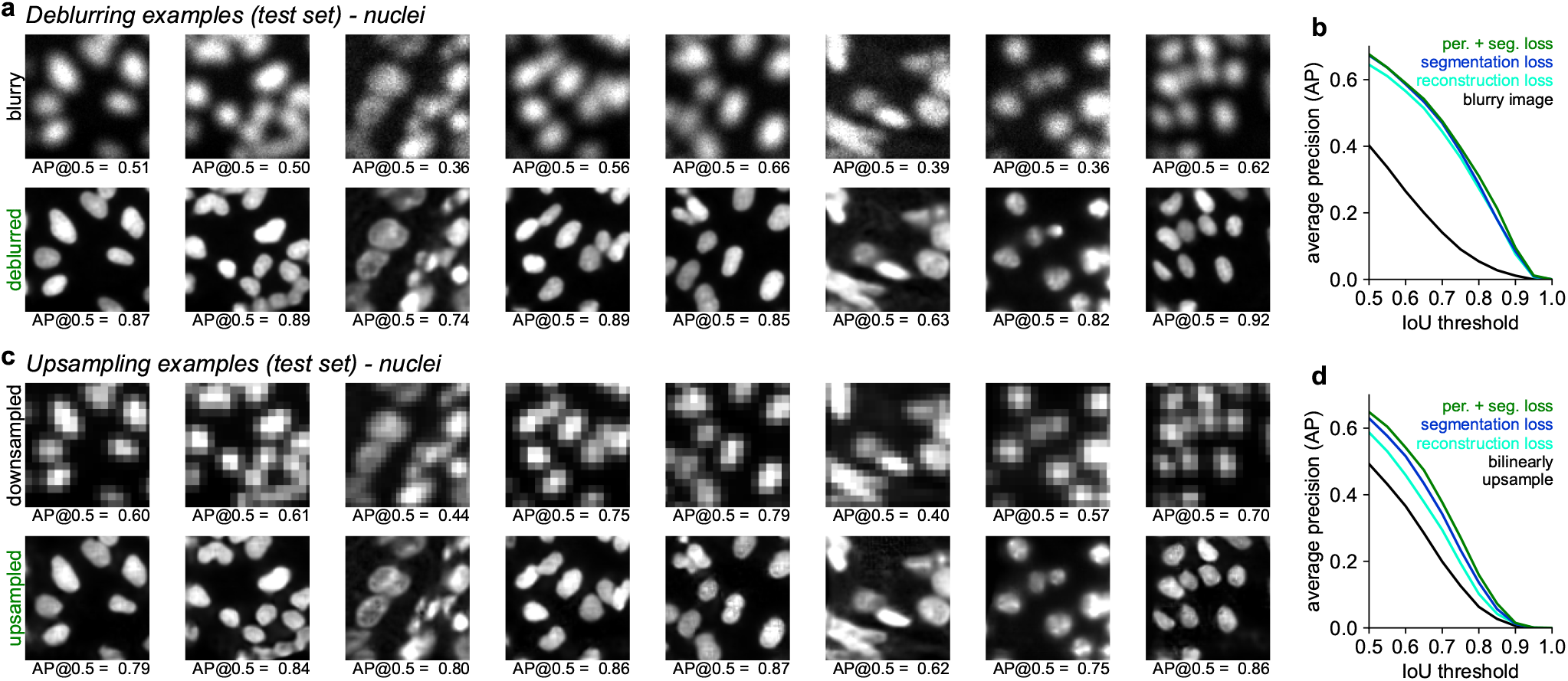
Deblurring and upsampling nuclear images. Same as Figure 4, for the Nuclei dataset.

**Figure S5:**
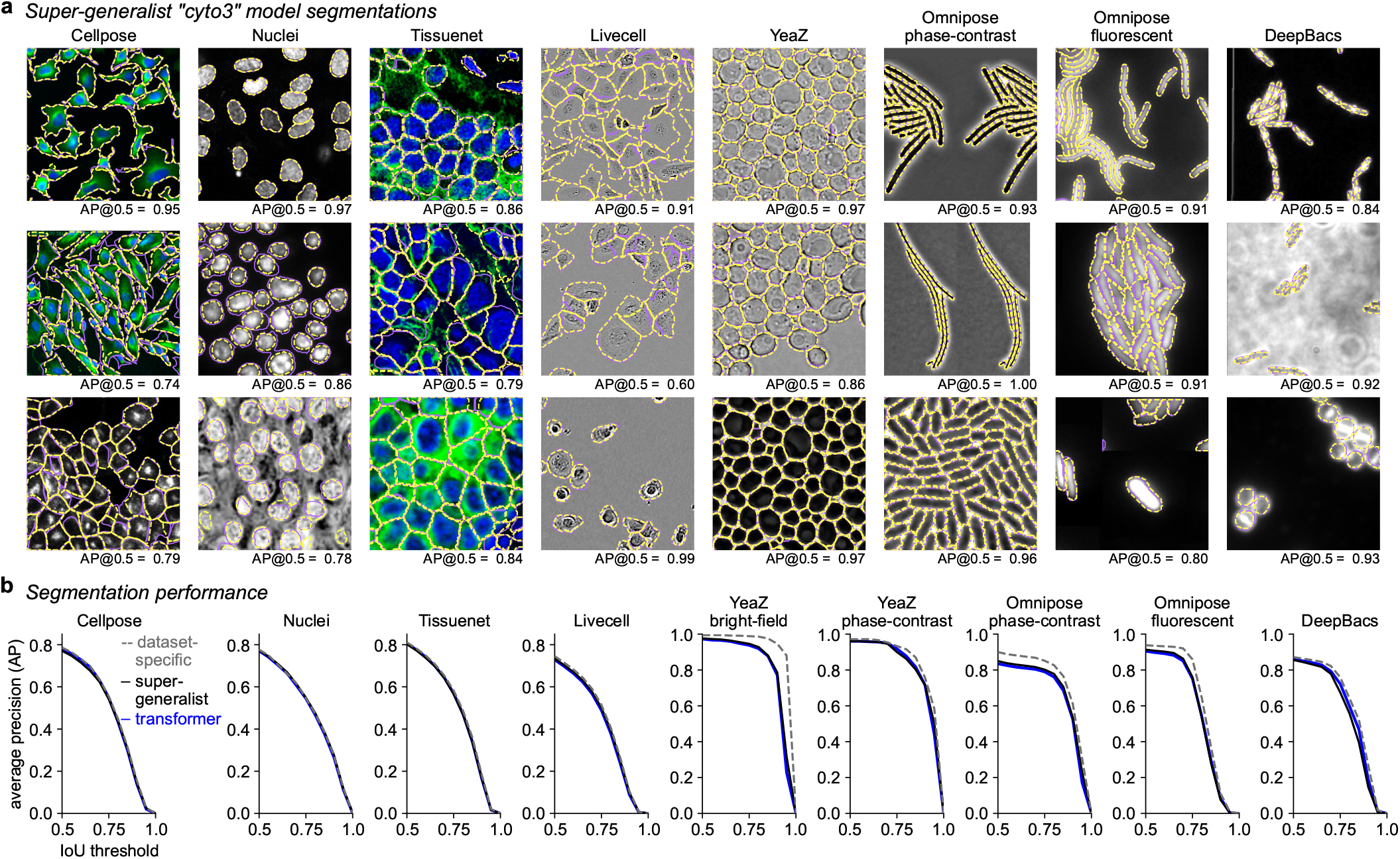
Segmentation performance of the ‘super-generalist’ model. A single model was trained to segment 9 datasets of cellular and nuclear images. **a**, Three example images from each test set (YeaZ bright-field and phase-contrast are combined), with ground-truth segmentations overlaid in purple, and super-generalist model segmentations in yellow. The AP@0.5 is the average precision of the segmentation at an IoU threshold of 0.5. **b**, Segmentation performance on the different test sets, averaged across all test images, of the super-generalist model, the dataset-specific models and the transformer model trained on all datasets.

